# Smooth Muscle Dysfunction Drives Cerebrovascular Reserve Failure and End-Organ Brain Injury

**DOI:** 10.64898/2026.01.26.701890

**Authors:** Takahiko Imai, Vijai Krishnan, James H Lai, Elyssa Alber, Lydia Hawley, Aarushi Gandhi, Pazhanichamy Kalailingam, Joanna Yang, Diana Tambala, David C Hike, Xiaoqing Alice Zhou, Claire Fong, Benjamin Ondeck, Emily T Da Cruz, Xiaochen Liu, Angelyna K Siv, Miran Öncel, Sabyasachi Das, Sava Sakadžić, Cenk Ayata, Xin Yu, Mark E. Lindsay, Patricia L Musolino, David Y Chung

## Abstract

**Background:** Failure of cerebrovascular reserve is a fundamental determinant of ischemic vulnerability, yet the mechanisms by which vascular smooth muscle dysfunction compromises reserve and predisposes the brain to injury remain incompletely defined. We therefore tested whether a pathogenic smooth muscle mutation produces a baseline failure of cerebrovascular reserve sufficient to render the brain vulnerable to hypoperfusion, even in the absence of fixed arterial occlusion.

**Methods:** We examined cerebrovascular structure, hemodynamics, and reserve in a genetically defined mouse model of ACTA2-associated multisystemic smooth muscle dysfunction syndrome with systemic or brain-restricted expression of the mutant allele. Cerebral artery morphology was assessed using magnetic resonance angiography and black ink angiography. Vascular smooth muscle phenotype was evaluated by immunohistochemistry and proliferation assays. Blood pressure reactivity and cerebral blood flow (CBF) were measured simultaneously using femoral arterial catheterization and laser speckle flowmetry during vasoactive challenges and controlled hypotension. Cerebrovascular stress responses were tested using unilateral common carotid artery occlusion. Downstream brain effects were assessed by histology, resting state functional connectivity imaging, and behavioral testing.

**Results:** Impaired smooth muscle contractility drove rectification and narrowing of major cerebral arteries, downregulation of contractile markers, and increased vascular cell proliferation. These structural changes produced a distinct physiological phenotype: mutant mice exhibited blunted vasoreactivity, diminished spontaneous vasodynamic activity, and a downward shift in the blood pressure-CBF relationship across a wide range of arterial pressures, consistent with loss of cerebrovascular reserve. As a result, CBF was reduced at baseline and could not be maintained during hypotension or acute vascular stress. During carotid occlusion, mutant mice showed impaired compensatory perfusion, greater physiological instability, and worse behavioral outcomes. Chronic reserve failure coincided with white matter loss, reduced neuronal density, disrupted large-scale functional connectivity, and deficits in locomotion, anxiety-related behavior, and working memory.

**Conclusions:** Pathogenic smooth muscle dysfunction caused by ACTA2 mutation produces a baseline failure of cerebrovascular reserve that renders the brain vulnerable to hypoperfusion and stress-induced ischemic injury. These findings establish cerebrovascular reserve failure as a central physiological mechanism linking vascular dysfunction to end-organ brain injury and identify reserve preservation as a critical, potentially actionable determinant of brain health in hypotension-prone vascular disease.

**Clinical Perspective:** *What Is New?:*

- ACTA2 smooth muscle dysfunction produces baseline cerebrovascular reserve impairment, with reduced cerebral blood flow and a downward-shifted pressure-flow relationship in the absence of critical large-vessel occlusion.
- Vascular tone dysregulation is coupled to end-organ brain injury, including white matter and neuronal loss, disrupted functional connectivity, and behavioral deficits.
- The results support complementary disease mechanisms in ACTA2 vasculopathy: baseline reserve limitation and injury-provoked occlusive remodeling.

*Clinical Implications:*

- Patients with ACTA2 vasculopathy may be vulnerable to ischemic brain injury during hypotension or systemic stress despite the absence of critical stenosis or occlusion on routine imaging.
- Peri-procedural and acute-care management should emphasize preserving perfusion pressure and cerebrovascular reserve (e.g., during anesthesia, dehydration, or systemic illness).
- More broadly, cerebrovascular reserve is a clinically relevant, potentially modifiable determinant of brain health in hypotension-prone vasculopathies and conditions characterized by impaired vascular reactivity.

## Introduction

Vascular smooth muscle cell (SMC) contractility is essential for arterial tone, blood pressure reactivity, and protection of end-organ perfusion through autoregulation and vascular reserve. These mechanisms allow the circulation to buffer physiologic stress, maintain stable tissue perfusion, and prevent ischemic injury during fluctuations in systemic pressure. When SMC function is impaired, the resulting loss of vascular reserve renders end organs vulnerable to hypoperfusion, particularly under conditions of stress such as hypotension, anesthesia, or acute vascular obstruction. Despite the central role of smooth muscle-dependent vascular reactivity in cardiovascular physiology, the consequences of isolated and baseline SMC dysfunction on end-organ injury remain incompletely defined, in part because most clinical and experimental models involve multifactorial vascular and parenchymal disease.

The brain represents a uniquely sensitive end organ in this context because of its high metabolic demand, limited ischemic tolerance, and reliance on tightly regulated cerebral blood flow to sustain distributed neural networks. Even modest or intermittent reductions in perfusion can preferentially injure long-range white matter tracts and disrupt functional connectivity in the absence of large territorial infarction. Consistent with this vulnerability, cerebrovascular small and large vessel disease are major contributors to vascular cognitive impairment and dementia (VCID), a progressive syndrome that reflects cumulative vascular injury rather than focal stroke alone.^1-4^ Common cardiovascular and aging-related conditions (including hypertension, diabetes, atherosclerosis, Alzheimer’s proteinopathy, cerebral amyloid angiopathy, and aging) are strongly associated with cerebral small vessel disease and impaired perfusion.^4^ Across these disorders, dysfunction of vascular SMCs and pericytes limits the dynamic regulation of cerebral blood flow (CBF), constraining cerebrovascular reserve and impairing the brain’s ability to tolerate physiologic stress. However, because vascular pathology in these settings typically coexists with neurodegenerative and inflammatory processes, the direct contribution of baseline smooth muscle dysfunction to cerebrovascular reserve failure and downstream brain injury has remained difficult to define.

In the present study, we leveraged a genetically defined mouse model of ACTA2 R179-mediated smooth muscle dysfunction to directly test how impaired SMC contractility affects cerebrovascular structure, hemodynamics, and reserve. The ACTA2^R179H/+^ allele is a flox-activated exon advancement model that enables tissue-specific expression of the mutant allele following Cre recombination.^5^ To isolate the contribution of vascular smooth muscle dysfunction to cerebrovascular physiology, we employed two complementary Cre-driver strategies. Systemic activation of the mutation in all SMCs and pericytes was achieved using the smooth muscle myosin heavy chain promoter (Myh11-Cre), generating systemic ACTA2 (sACTA2) mice.^5^ In parallel, brain-restricted expression within the anterior circulation was achieved using a cardiac neural crest–specific Cre (Wnt1-Cre), generating brain ACTA2 (bACTA2) mice. Using these models, we examined cerebral artery structure, blood pressure reactivity, and static and dynamic cerebral blood flow to define cerebrovascular reserve failure under baseline and stress conditions. We further assessed the downstream consequences of cerebrovascular reserve failure on white matter integrity, functional connectivity, and behavior as end-organ manifestations of vascular dysfunction.

## Methods

### Animal Models and Study Approval

All procedures involving animals were approved by the Mass General Brigham Institutional Animal Care and Use Committee (Protocols 2012N000196 and 2018N000067) and conducted in accordance with the ARRIVE 2.0 guidelines. Mice were housed in specific-pathogen-free conditions on a 12-hour light/dark cycle with ad libitum access to food and water. Environmental enrichment was provided in the form of nesting material and igloos. Body weight, behavior, and general health were monitored throughout the study.

#### Study design, experimental unit, randomization, blinding, and sample size

Mice were assigned to genotype-defined experimental groups (control Acta2^R179Hfl/+^ vs Cre-activated Acta2^R179Hfl/+^ as indicated for each experiment). The experimental unit was a single mouse for all *in vivo* experiments. For *ex vivo*/cell-based assays, the experimental unit was a culture well. Because group assignment was determined by genotype, randomization to genotype was not applicable. Where feasible, experimental run order was counterbalanced across genotypes; perfect counterbalancing was not always achievable due to variable mortality and availability of ACTA2 mutant mice prior to study procedures. Investigators were blinded to genotype during conduct, outcome assessment, and data analysis whenever feasible; genotype was revealed only after data processing/quantification was complete. Sample sizes for each experiment are reported in the corresponding figure legends and tables. No *a priori* power calculation was performed; sample sizes were based on prior experience with these assays and feasibility constraints.

#### Inclusion/exclusion criteria

Exclusion criteria were defined *a priori* as technical failure of key procedures (e.g., unsuccessful femoral catheterization, unstable anesthesia/physiology precluding completion of protocol, major motion artifact or incomplete MRI acquisition, inability to obtain interpretable laser speckle/optical signals). No exclusions were made other than instances of experimental failure as indicated in the corresponding figure legends and tables. To minimize confounding, age and sex were recorded for each cohort and evaluated in post hoc analyses (e.g., age/sex effects or adjustment), as indicated.

The conditional *Acta2^R179Hfl/+^* mouse model used in this study was previously described.^5^ This mouse contains a conditional allele in which Cre-recombinase activation of alternative exons replaces arginine 179 with histidine. To generate mice expressing the mutant allele in smooth muscle cells (sACTA2), female *Acta2^R179Hfl/fl^* mice were crossed with male B6.Cg-Tg(Myh11-cre,-EGFP)2Mik/J mice (JAX #007742, The Jackson Laboratory, Bar Harbor, ME) to generate *Myh11-cre:Acta2^R179Hfl/+^* (sACTA2) and *Acta2^R179Hfl/+^* mice (control). For brain-specific expression (bACTA2), female *Acta2^R179Hfl/fl^* mice were crossed with male 129S4.Cg-*E2f1Tg(Wnt1-Cre)2Sor*/J mice (JAX #022137) to generate *Wnt1-cre:Acta2^R179Hfl/+^* (bACTA2) and *Acta2^R179Hfl/+^* mice (control). All mice were maintained on a C57BL/6J background. Analysis of smooth muscle cells from activated *Acta2^R179Hfl/+^* mice demonstrated 50% mutant allele activation.^5^ Animal characteristics (age, weight, and sex) for each experimental cohort are summarized in Tables 1–7.

**Table 1.**
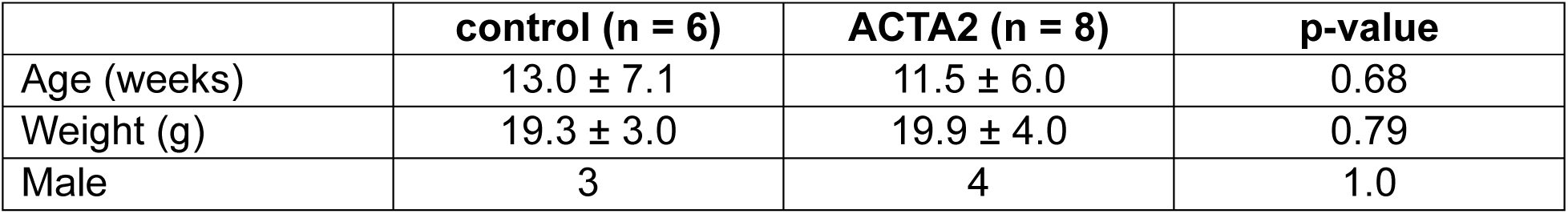
Age and weights of MRI Fig 2A.

**Table 2.** Black ink (Fig 2B and 3A)

|  | control (n = 12) | ACTA2 (n = 10) | p-value |
| --- | --- | --- | --- |
| Age (weeks) | 11.3 ± 2.2 | 11.5 ± 2.3 | 0.88 |
| Weight (g) | 25.3 ± 2.5 | 21.2 ± 6.2 | 0.06 |
| Male | 9 | 5 | 0.38 |

**Table 3.** Age, weight, and arterial blood gas values for SNP-PE experiments. Mean ± SD.

|  | control (n = 8) | ACTA2 (n = 7) | p-value |
| --- | --- | --- | --- |
| Age (weeks) | 7.8 ± 0.7 | 7.8 ± 0.5 | 0.96 |
| Weight (g) | 23.4 ± 1.4 | 19.5 ± 3.5 | 0.03 |
| Male | 8 | 4 | 0.077 |
| pH | 7.28 ± 0.07 | 7.28 ± 0.05 | 0.98 |
| dpCO<sub>2</sub> | 47 ± 11 | 51 ± 11 | 0.48 |
| pO<sub>2</sub> | 118 ± 34 | 133 ± 25 | 0.34 |

**Table 4.** Exsanguination experiments (Speckle Fig 2).

|  | control (n = 10) | ACTA2 (n = 10) | p-value |
| --- | --- | --- | --- |
| Age (weeks) | 11.0 ± 1.8 | 11.5 ± 2.3 | 0.63 |
| Weight (g) | 24.7 ± 2.0 | 21.2 ± 6.2 | 0.12 |
| Male | 8 | 5 | 0.35 |
| pH | 7.2 ± 0.1 | 7.3 ± 0.1 | 0.20 |
| pCO<sub>2</sub> | 59.1 ± 20.1 | 58.8 ± 14.5 | 0.98 |
| pO<sub>2</sub> | 113.7 ± 19.8 | 112.2 ± 36.9 | 0.92 |

**Table 5.** Common carotid artery occlusion (CCAO) experimental table.

|  | control (n =14) | ACTA2 (n = 18) | p-value |
| --- | --- | --- | --- |
| Age (weeks) | 7.9±0.7 | 8.4±1.0 | 0.18 |
| Weight (g) | 21.0±3.4 | 21.1±4.7 | 0.91 |
| Surgical duration | 19.9±8.4 | 18.6±2.9 | 0.52 |
| Male | 7 | 11 | 0.72 |

**Table 6.**
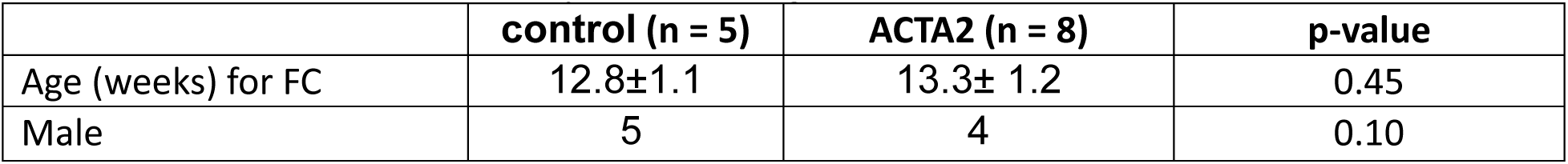
Functional connectivity table for Figure 8.

**Table 7.** Behavior table for Figure 8.

| <u>Behavior</u> | **control (n = 6)** | **ACTA2 (n = 9)** | **p-value** |
| --- | --- | --- | --- |
| Age (weeks) for behavior | 12.6±1.1 | 13.1±1.0 | 0.41 |
| Male | 5 | 4 | 0.29 |

#### Human MRI

The human neuroimaging data presented in this manuscript were acquired as part of an Institutional Review Board-approved Retrospective and Prospective Natural History of Genetic Vasculopathies study at Massachusetts General Hospital (Protocol 2023P000821). Written informed consent was obtained from parents or legal guardians, with assent from participants when appropriate. Magnetic resonance imaging (MRI) and magnetic resonance angiography (MRA) were performed on clinically indicated scanners using standardized pediatric and adult neurovascular imaging protocols. MRI sequences included diffusion-weighted imaging (DWI) with corresponding apparent diffusion coefficient (ADC) maps for detection of acute ischemia, and fluid-attenuated inversion recovery (FLAIR) for assessment of chronic white matter injury. Time-of-flight MRA was used to evaluate intracranial arterial anatomy and identify stenosis or occlusion of major cerebral vessels. Imaging was obtained at acute and longitudinal time points, as clinically indicated, allowing evaluation of lesion evolution, penumbral expansion, and long-term structural sequelae. Images were reviewed independently by study investigators with expertise in pediatric stroke and cerebrovascular disease. Lesions were categorized based on anatomical distribution (deep white matter watershed, arterial territory infarction), temporal evolution, and associated vascular findings on MRA.

### Mouse magnetic resonance image acquisition and processing

Time-of-flight (TOF) MRA was performed on ACTA2 and control mice under isoflurane anesthesia to assess vessel morphology. For scanning, animals were anesthetized using isoflurane. Induction was carried out under 5% isoflurane in 0.8 L/min of medical air supplemented by 0.2 L/min of O₂, and after the animals were secured in an MR-compatible holder using ear and tooth bars, isoflurane was reduced and maintained at 1.5%. Fine adjustments were made to ensure the animals remained stable during their time inside the magnet. The monitoring of the animals’ physiological status was facilitated using a SAII Small Animal Monitoring and Gating System (Model 1030, SAII, USA) tracking respiration and body temperature. The respiration rate was constantly observed with a pressure-sensitive pad and kept within 90–120 breaths per minute. To maintain the animals at a steady temperature of 37°C, warm air was blown over the animal while in the MR scanner’s bore and verified using a rectal probe thermometer. All imaging was performed using a horizontal 14 Tesla Magnex magnet (Magnex Scientific, UK), interfaced with a Bruker BioSpin AV Neo console and equipped with a high-performance gradient insert (Resonance Research Inc., MA, USA) capable of delivering a peak gradient strength of 1.2 T/m. An implantable RF transceiver surface coil with an outer diameter of 9.5 mm (Mribot Inc.) was used for image acquisition. To achieve high spatial resolution necessary for detailed vascular imaging, we employed a 3D Fast Low Angle Shot (FLASH) with an isotropic resolution of 100 μm. Acquisition parameters were as follows: repetition time (TR) = 1000 ms, echo time (TE) = 2.7 ms, matrix size = 128 × 60 × 90, and bandwidth = 79,365 Hz. The total acquisition time for each scan was approximately 1 hour and 9 minutes (1.15 h). This protocol enabled high-quality visualization of fine microvascular structures in vivo. Image reconstruction and subsequent processing were performed using the AFNI (Analysis of Functional NeuroImages) software suite. Raw Bruker 2dseq data were converted into AFNI format using the to3d function with anatomical field-of-view orientation and outlier skipping enabled. To standardize voxel intensities across datasets, images were normalized using a min–max scaling approach. Minimum and maximum signal values were extracted using 3dBrickStat, and intensity values were scaled to a 0–1 range using 3dcalc. All analyses were performed blinded to genotype.

### Black Ink Angiography and Circle of Willis Vessel Cross-Sectional Area

For *ex vivo* angiography, mice were transcardially perfused with 10-50 μL black India ink (Higgins, Chartpak Inc., Leeds, MA, USA), and brains were imaged under a dissecting microscope. Arterial morphology and curvature were assessed qualitatively. Diameters of the middle cerebral artery (MCA), anterior cerebral artery (ACA), internal cerebral artery (ICA), first segment of the posterior cerebral artery (P1), posterior communicating artery (pcomm), and basilar arteries were measured from brain images obtained after black ink perfusion using FIJI/ImageJ software (v1.54f, NIH, Bethesda, MD, USA). Cross-sectional vascular areas were calculated by averaging bilateral vessel diameters. All analyses were performed blinded to genotype.

### Cerebral VSMCs Isolation Protocol

Brains from control and Acta2 mutant mice were removed after perfusion with cold PBS and placed in ice-cold DPBS containing 1% penicillin–streptomycin. Cerebral arteries, including the circle of Willis and its branches, were dissected under a stereomicroscope, rinsed with DPBS, and digested with collagenase type II (1 mg/mL) at 37°C for 15 min with shaking. Digestion was terminated by adding DMEM/F12 supplemented with 10% FBS, followed by centrifugation at 1,500 rpm for 5 min. The pellet was resuspended in DMEM/F12 containing 20% FBS and 1% penicillin–streptomycin and plated onto 0.1% gelatin–coated culture dishes. Cells were incubated at 37°C in a humidified 5% CO₂ atmosphere, and tissue and cell attachment was assessed at 48–72 h, with medium changes performed as needed.

### Smooth Muscle Cell Proliferation Assay

Primary cerebral vascular smooth muscle cells (VSMCs) were isolated and plated at 5,000 cells per well in black-walled 96-well plates (Corning, Corning, NY, USA). After 5 h attachment at 37°C and 5% CO₂, media was replaced with complete growth medium containing NucSpot® 647 nuclear stain (1:2000 dilution; Biotium, Fremont, CA; Cat. #40082). Plates were imaged every 2 h for 72 h using an Incucyte® live-cell imaging system (Sartorius, Göttingen, Germany). Nuclei were quantified with Incucyte® software.

### Tissue Collection, Immunofluorescence, and histology

For histology and immunohistochemistry experiments, mice were anesthetized and transcardially perfused with phosphate-buffered saline (PBS) followed by 4% paraformaldehyde (PFA) in PBS. Brains were post-fixed overnight in 4% PFA at 4 °C, cryoprotected in 30% sucrose, and embedded in optimal cutting temperature (OCT) compound. Coronal sections (8 μm thick) were prepared using a cryostat and mounted on Superfrost Plus slides (Thermo Fisher Scientific, Waltham, MA, USA). Sections were permeabilized with 0.3% Triton X-100 in PBS and blocked with 3% bovine serum albumin (BSA) for 1 hour at room temperature.

For cerebral VSMC F-actin visualization, cells were stained using ActinGreen 488 ReadyProbes Reagent (Thermo Fisher Scientific) following fixation and permeabilization. After incubation with primary antibodies and fluorophore-conjugated secondary antibodies for smooth muscle actin (SMA), cells were incubated with ActinGreen 488 (two drops per mL in PBS) for 30 minutes at room temperature, protected from light. Sections were then washed three times with PBS, counterstained with DAPI, prior to imaging. For other immunohistochemistry experiments, primary antibodies were applied overnight at 4 °C in 1% BSA/PBS, as follows: Anti-alpha smooth muscle actin (α-SMA, Abcam, AB7817) and secondary Alexa Fluor 647 goat anti-mouse (Abcam, AB150115). Anti-smoothelin (Santa Cruz Biotechnology, SC-376902) and secondary Alexa Fluor 647 goat anti-mouse (Abcam, AB150115). Anti-transgelin (Santa Cruz Biotechnology, SC-53932) and secondary Alexa Fluor 647 goat anti-mouse (Abcam, AB150115). Anti-calponin (Abcam, AB46794) and secondary Alexa Fluor 750 goat anti-rabbit (Invitrogen, 2831371). Anti-NeuN (Abcam, AB177487) and secondary Alexa Fluor 750 goat anti-rabbit (Invitrogen, 2831371). Anti–myelin basic protein (MBP, Invitrogen, PA1-46447) and secondary: Alexa Fluor 750 goat anti-rabbit (Invitrogen, 2831371). Following three PBS washes, sections were incubated for 1 hour at room temperature with fluorophore-conjugated secondary antibodies and counterstained with DAPI (1 μg/mL; Thermo Fisher Scientific, Cat. #D1306). Slides were mounted with Fluoromount-G (SouthernBiotech, Birmingham, AL) and imaged using a Zeiss LSM 800 confocal microscope (Carl Zeiss AG, Oberkochen, Germany).

Images were analyzed using FIJI/ImageJ software. Regions of interest (ROIs) were manually drawn for cortical vessels, motor cortex, corpus callosum, and striatum. For contractile markers (SMA, smoothelin, transgelin, calponin), mean fluorescence intensity was measured within each vessel ROI. NeuN-positive cells were counted using the “Analyze Particles” function following thresholding and were normalized to area (cells/mm²). MBP signal intensity was quantified as mean gray value within white matter ROIs.

### Blood Pressure Measurement and Pharmacological Response

Mice were anesthetized with isoflurane (3% induction, 1.5–2.0% maintenance), and concurrent femoral artery and vein catheterization was performed using PE-10 tubing (BD Intramedic™, Becton Dickinson, Franklin Lakes, NJ, USA). Mean arterial pressure (MAP) was recorded using a PowerLab/16SP system (ADInstruments, Colorado Springs, CO, USA) with a pressure transducer (Edwards Lifesciences, Irvine, CA, USA).

Mice were transitioned to tribromoethanol anesthesia (Avertin, 250–400 mg/kg IP in 1.25% tert-amyl alcohol; Sigma-Aldrich, St. Louis, MO, USA). Sodium nitroprusside (SNP, Sigma-Aldrich, 0.2 mg SNP per 1 g mouse weight in 1 mL saline, 71778, St. Louis, MO, USA) and phenylephrine (PE, Sigma-Aldrich, 0.2 mg/mL, P1240000, St. Louis, MO, USA) were infused via a femoral vein using a syringe pump (KD Scientific, Legato 100 Infuse Single Syrine, 788100, Holliston, MA, USA). Dose-response MAP was normalized to the timepoint between SNP discontinuation and PE initiation. Arterial blood gases were analyzed pre- and post-intervention using the OPTI CCA-TS2 analyzer (OPTI Medical Systems, Roswell, GA, USA).

#### Laser Speckle Cerebral Blood Flow (CBF)

Relative CBF was recorded in sACTA, bACTA2, and control mice using custom-built laser speckle flowmetry setups^6^ in 2 distinct experiments. Common to both experiments, a femoral artery catheter enabled simultaneous MAP measurement. After induction and maintenance with isoflurane, the scalp was incised taking care to immediately apply mineral oil to the skull surface. The skull was further cleared of periosteum while applying mineral oil to maintain transparency through the bone. After several minutes, excess oil was removed from the surface of the skull to minimize light reflections. Diffuse laser light was directed onto the skull surface (785 nm laser diode, L785P090 Thorlabs, Newton, NJ, USA). Time-domain speckle contrast images were calculated by taking the mean and standard deviation of 50 images. The speckle contrast images were converted to 1/tc maps which are proportional to CBF. Whole-brain ROIs were analyzed using custom MATLAB scripts for MAP-CBF curve generation. For the SNP/PE experiments (**Figure 5a-f**), the frequency of 1/tc map acquisition was 1 Hz. To ensure comparability across experimental cohorts, 1/tc was normalized to account for its sensitivity to subtle changes in the experimental setup over time (e.g., laser light intensity). A separate experiment was carried out where LSF CBF imaging was determined during stepwise blood withdrawal (25-100 µL every 10 min) from a femoral artery catheter (**Figure 5 g-h**). Imaging continued until circulatory collapse. For this experiment, mice were measured within a narrow time window, which minimized drift in experimental conditions. As a result, raw 1/tc values were directly comparable across animals and were used without normalization.

#### Common Carotid Artery Occlusion (CCAO)

Unilateral common carotid artery occlusion (CCAO) was performed to assess the ability of mice to maintain cerebral blood flow (CBF) under acute cerebrovascular stress. Control and sACTA2 (ACTA2 R179H) mice were studied (n = 14 control, n = 18 sACTA2), as shown in the experimental timeline (**Figure 6a**) and **Table 5**. Mice were anesthetized with isoflurane (induction: 3%, maintenance: 1.5-2.0%) delivered in 70%N_2_ and 30% O_2_, and body temperature was maintained at 36–37°C using a feedback-controlled heating pad (FHC, Bowdoinham, ME, USA). Ophthalmic ointment was applied to prevent corneal drying.

A midline cervical incision was made under aseptic conditions, and the left common carotid artery (CCA) was carefully exposed by blunt dissection, avoiding damage to the vagus nerve. Permanent occlusion was achieved using 6-0 silk suture placed around the artery and secured with a knot. Successful occlusion was confirmed by an immediate reduction in ipsilateral cerebral blood flow measured by laser Doppler flowmetry (see below). The incision was closed with sutures, and animals were allowed to recover under a warming lamp before return to their home cage. Relative cerebral blood flow was measured using laser Doppler flowmetry (Perimed, Ardmore, PA, USA).

After induction of anesthesia and prior to carotid ligation, a laser Doppler probe was positioned and secured over the left middle cerebral artery (MCA) territory with the skull intact. Baseline CBF was recorded prior to ligation. Following permanent ligation of the left CCA, CBF was continuously monitored for 10 minutes post-occlusion. Residual CBF was calculated as the mean post-occlusion signal expressed as a percentage of each animal’s baseline value. Animals were monitored daily for 14 days post-CCAO. Body weight was recorded daily and expressed as percent change from pre-operative baseline. Neurological testing was performed with a 5-point score as described below. Survival was recorded daily up to 14 days post-CCAO. At Day 14, surviving animals were euthanized and perfused, as indicated in Figure 6a.

#### Peri-procedural animal welfare and humane endpoints

All surgical procedures were performed under aseptic conditions with measures to minimize pain and distress, including maintenance of normothermia, ocular lubrication, and post-operative monitoring. Animals were assessed at least daily for weight loss, hydration status, activity, grooming, and neurological deficits. Humane endpoints included criteria specified in the IACUC protocol (e.g., severe neurological impairment, inability to ambulate or access food/water, moribund appearance, or excessive weight loss), and animals meeting endpoint criteria were euthanized according to approved protocols.

### CBV Fluctuation and Functional Connectivity Imaging

To evaluate resting vasodynamics, optical intrinsic signal imaging was performed through a chronic glass coverslip over the intact skull as previously described.^7,8^ In short, mice were anesthetized with 5% isoflurane with a combination of 70:30 N_2_O_2_:O_2_ and maintained at 1.5%-2.0% while breathing spontaneously. Once fully anesthetized, the head was fixed in a stereotactic frame. Eye lubricant was applied, and the body temperature was maintained at (37.0 ± 0.1°C) using a rectal probe and heating pad. The fur was split along the midline, and an incision was made in the scalp along the dorsal skull surface. Once the periosteum was removed, and the scalp reflected outward, gauze was placed inside, between the skin and the skull, to help dry out the scalp. A glass coverslip (12 mm diameter, Electron Microscopy Sciences, Cat #72196-12, Hatfield, PA, USA) was trimmed with a diamond pen to reflect the shape of the dorsal skull surface. The coverslip was then adhered to the skull surface with C&B Metabond cement, which was mixed in a pre-chilled ceramic mixing dish using six drops of Quick Base, one drop of catalyst, and one scoop of Clear L-Powder (Product numbers S398, S371, and S399, Parkell, Edgewood, NY, USA) as previously described.^7^ Mice were allowed to recover for at least 4 days. For optical imaging experiments, mice were anesthetized using 2,2,2-tribromoethanol (TBE), also known as Avertin, which enables excellent RSFC signals.^9^ Injectable Avertin was prepared by combining 0.5mL of stock solution (10g TBE in 6.25mL tert-amyl alcohol; T48402-25G, Sigma-Aldrich, St. Louis MO, USA and P005925M, Fisher Scientific, Peabody, MA, USA, respectively) with 39.5 mL of 0.9% normal saline, protected from light, and stored at 4°C for no longer than one month. To smoothly induce the mouse to an adequate level of anesthesia where the withdrawal to a toe pinch was suppressed, TBE was injected intraperitoneally over two to three doses (0.3–0.4 mL initial dose, 0.1 mL at approximately 8 min, and 0.1 mL at approximately 12 min). More TBE was injected if the mouse still reacted to a toe pinch.

The animal was head-fixed in a stereotaxic frame, eye lubricant was applied, and the body temperature was maintained at (37.0 ± 0.1°C) using a rectal probe and heating pad. The previously implanted glass coverslip was cleaned using 70% EtOH and a cotton-tipped applicator. Imaging began at 18-20 minutes post initial dose of TBE. Three strobing red, blue, and green LEDs (LEDD1B T-Cube Driver, M625L3, M470L3, and M565L3 respectively, ThorLabs, Newton, NJ, USA) and a Basler camera (a2a 1920-150um Pro, Basler AG, Germany) were positioned over the anesthetized mouse.

Images were collected for 12 minutes using Pylon viewer software (Basler AG, Germany), at 30 frames per second (fps). The raw reflectance data were analyzed using custom MATLAB scripts. The modified Beer-Lambert law was used to calculate total (HbT), oxy- (HbO), and deoxy-hemoglobin (HbR) maps.^10^ The power spectral density of resting state oscillations of total hemoglobin (HbT) was quantified at different animal ages with and without global signal regression. Longitudinal resting state functional connectivity (RSFC) analysis was also performed on the processed HbT, HbO, and HbR data. An interhemispheric bihemispheric connectivity index (BCI) was calculated based on the correlation of each pixel with its homotopic counterpart in the contralateral hemisphere. A global connectivity index (GCI) was calculated based on the positive correlations of a pixel with all other pixels across the dorsal surface of the brain.^7,9^ All correlations were Fisher z-transformed so that mean and standard deviations could be determined for statistical comparisons across genotypes and age.

#### Behavioral Testing

For CCAO experiments, neurological function was assessed using a 5-point neurological deficit score, performed by an investigator blinded to genotype, as follows: 0, normal; 1, turning the torso to one side when held by the tail; 2, circling behavior; 3, cannot bear weight on 1 side; and 4, no spontaneous movement or barrel rolling).^11^ Otherwise, neurological function was scored using a 14-point Modified neurologic severity score (mNSS) assessing motor function, reflexes, and coordination.^12^ The open field test (OFT) was captured with a Logitech USB web.

Camera and performed in a 17x13 in^2^ arena, and ambulation and thigmotaxis were quantified over 5 minutes using video tracking software via University of Pennsylvania automated phenotyping of behavior MATLAB code. (https://www.seas.upenn.edu/~molneuro/software.html). The Y-maze test was conducted for 8 minutes; number of arm entries and spontaneous alternations were manually tabulated.

#### Statistical Analysis and Illustration

All analyses were performed using Prism (GraphPad, San Diego, CA) and JMP (SAS Institute, Cary, NC). Data were assessed for normality. Between-group comparisons were made using unpaired t-tests or two-way ANOVA as appropriate. Longitudinal data were analyzed using mixed-effects models. p < 0.05 was considered significant. Error bars represent standard deviation (SD) unless otherwise indicated. Normality was assessed using standard graphical and/or formal tests within Prism/JMP; when distributional assumptions were not met, nonparametric or variance-robust approaches were used as appropriate. All statistical details are included in figure legends. Fisher’s Exact test was used to assess sex distribution. Schematic representations were generated using BioRender to illustrate hypothesized mechanisms of ischemic injury related to impaired cerebrovascular autoregulation in ACTA2 R179–associated syndrome (MSMDS).

#### Outcome measures

Outcome measures and sample sizes for each experiment are reported in the corresponding figure legends and tables. The primary outcome measure for hypothesis-testing was the MAP-CBF relationship (pressure-flow relationship) assessed during controlled hypotension/exsanguination; other measures were considered secondary.

#### Protocol registration and data availability

No prospective protocol registration was performed for this study. Data supporting the findings of this study are available from the corresponding author upon reasonable request.

## Results

ACTA2 R179H-associated multisystemic smooth muscle dysfunction syndrome (MSMDS) is characterized clinically by marked vulnerability to hypotension-associated ischemic injury. Affected patients develop deep watershed white matter lesions and propagation of injury across arterial territories during periods of systemic stress, often in the absence of fixed large-vessel occlusion (**Figure 1**).^13^ These injury patterns are not readily explained by focal arterial obstruction alone and instead suggest impaired autoregulation and limited cerebrovascular reserve as dominant mechanisms of ischemic vulnerability. We therefore examined vascular structural and physiological features that could account for this phenotype in a genetically defined mouse model of ACTA2-associated MSMDS.

**Figure 1.**
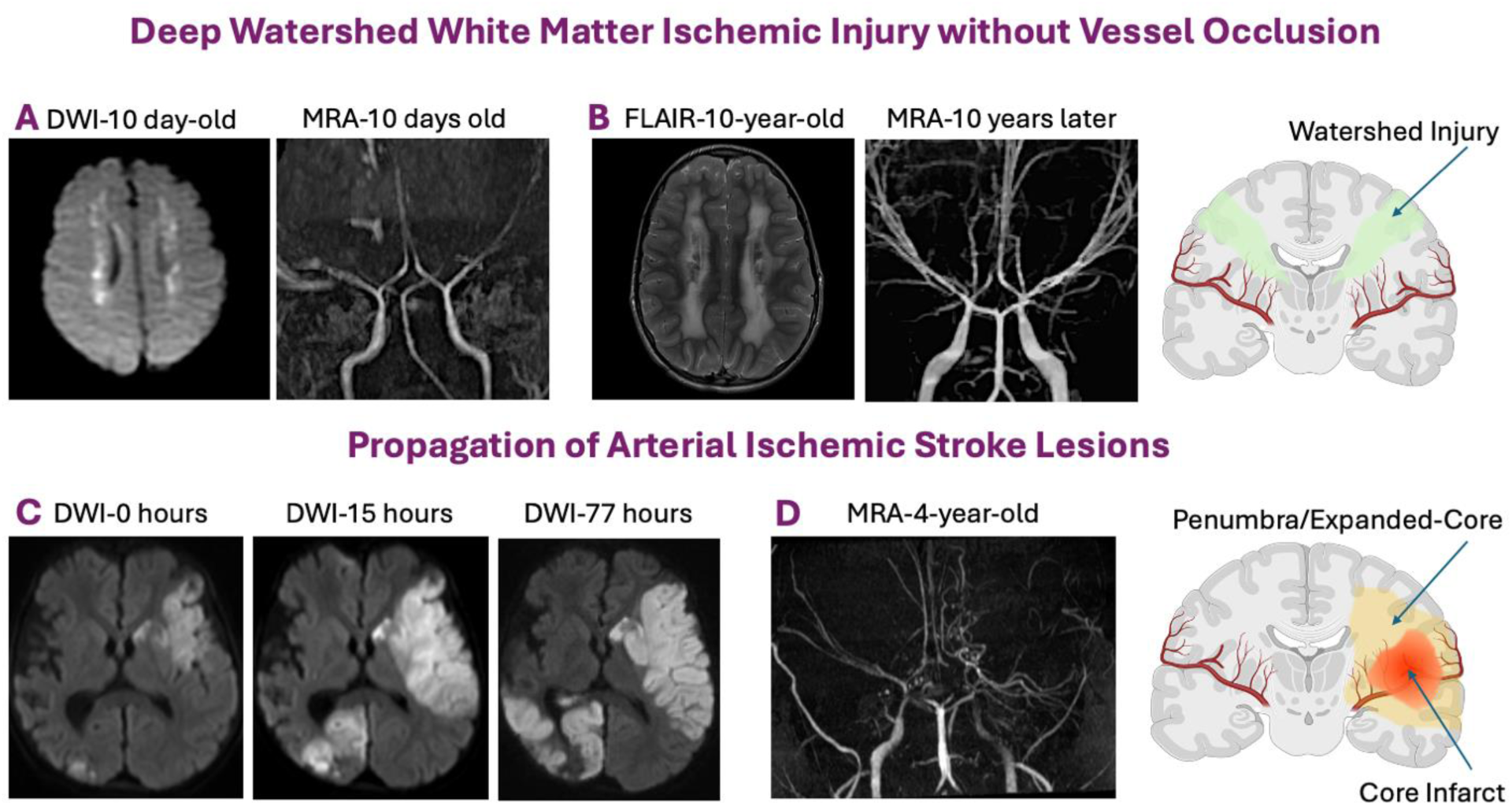
Patterns of brain injury in MSMDS are consistent with failure of cerebrovascular autoregulation and reserve. MRI and MR Angiograph images of ACTA2 R179 patients (A) Acute Deep White Matter Watershed Ischemic Injury demonstrated by hyperintensities on MRI Diffusion-Weighted Imaging (DWI) on a 10-day-old male 3 days after PDA surgical repair. MRA showed no evidence of cerebral arteries critical stenosis or occlusions suggesting hypoperfusion/hypoxic injury. (B) Long-term follow-up 10 years later shows chronic white-matter lesion as depicted by FLAIR hyperintensities largely corresponding with perinatal ischemic lesions and no cortical involvement despite significant progression of cerebral artery stenosis on MRA. (C) DWI Acute Arterial Ischemic Strokes lesions in a 5-year-old female and its expansion into contiguous tissue at risk (penumbra) over first 3 days. (D) MRA showed new L ICA occlusion, critical stenosis of R ICA, ACA and MCA without occlusion. Schematics of types of ischemic injury in ACTA2 R179.

To address this question, we studied a conditional Acta2^R179Hfl/+^ mouse model enabling smooth muscle–specific expression of the mutant allele.^5^ Systemic expression in vascular smooth muscle cells and pericytes was achieved using Myh11-Cre (sACTA2), whereas brain-restricted expression within the anterior circulation was achieved using Wnt1-Cre (bACTA2). These complementary models enabled interrogation of cerebrovascular structure, vascular reactivity, cerebral blood flow, and reserve under baseline and stress conditions.

ACTA2 R179H MSMDS is associated with a distinctive cerebrovascular anatomy characterized by rectification of the cerebral arterial system.^13^ Using *in vivo* non-contrast MRA—an approach analogous to clinical imaging—we observed reduced arterial curvature in the anterior cerebral and middle cerebral arteries of sACTA2 mice (n = 8) compared with controls (n = 6) (**Figure 2a**). Qualitative alterations in surface MCA geometry were confirmed in an independent cohort following black ink perfusion (sACTA2 n = 10; control n = 12) (**Figure 2b**).

**Figure 2.**
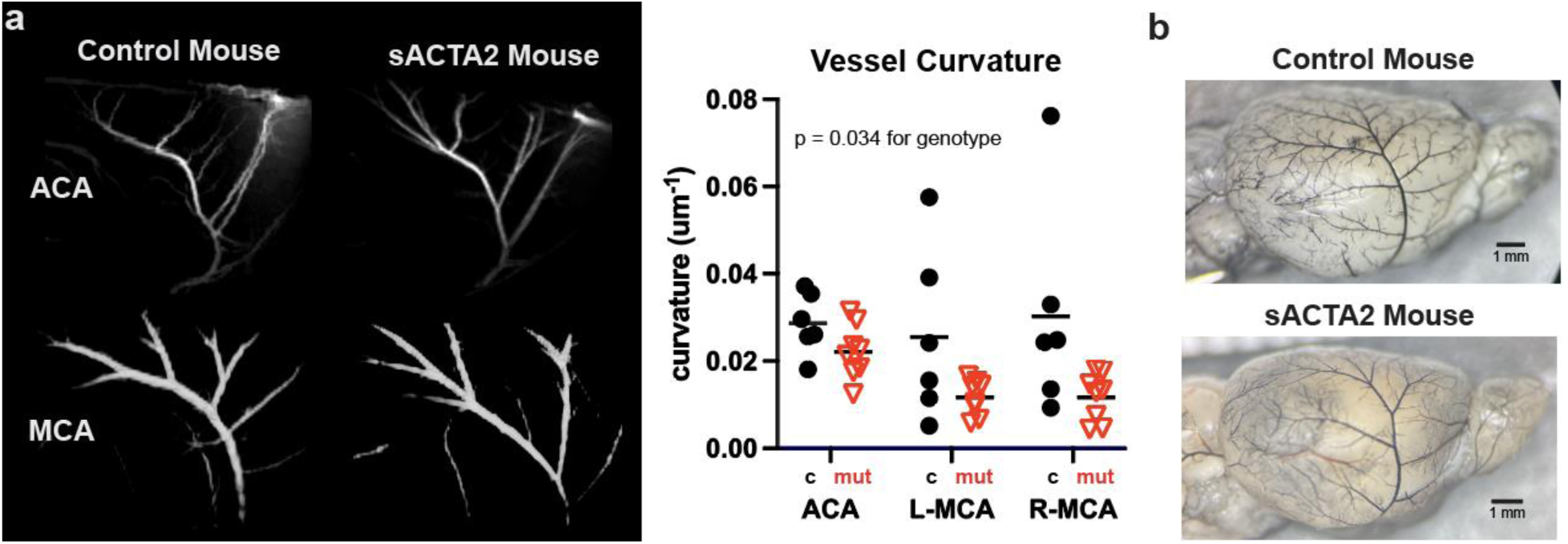
ACTA2 mice have cerebral artery rectification. (a) The ACTA2 mutation leads to rectification (straightening) of the anterior cerebral artery (ACA) and middle cerebral artery (MCA). Magnetic resonance angiograph (MRA) showing rectification of the ACA and MCA in sACTA2 mice that have systemic and cerebral expression of the mutation (n = 6 control, n = 8 sACTA2/mut). (b) Morphology of the MCAs visualized after black ink injection from control and sACTA2 brains (representative brains from n = 12 control, n = 10 sACTA2).

In the black ink cohort, diameters of proximal arteries at the skull base were measured and cross-sectional areas calculated. Mutant mice exhibited consistent reductions in arterial cross-sectional area compared with controls (**Figure 3a**), mirroring the human phenotype.^13^ Primary cultured vascular smooth muscle cells from sACTA2 mice demonstrated increased proliferation (**Figure 3b**), suggesting that luminal narrowing may be driven in part by smooth muscle hyperplasia, consistent with prior histopathologic observations in ACTA2 R179H cerebrovascular disease.^14^ Cytoskeletal organization was also perturbed: F-actin and smooth muscle actin (SMA) staining revealed distorted stress-fiber architecture and abnormal SMA localization in sACTA2 cells compared with controls (**Figure 3c**), consistent with previously reported ACTA2-associated cytoskeletal defects.^5,15^

**Figure 3.**
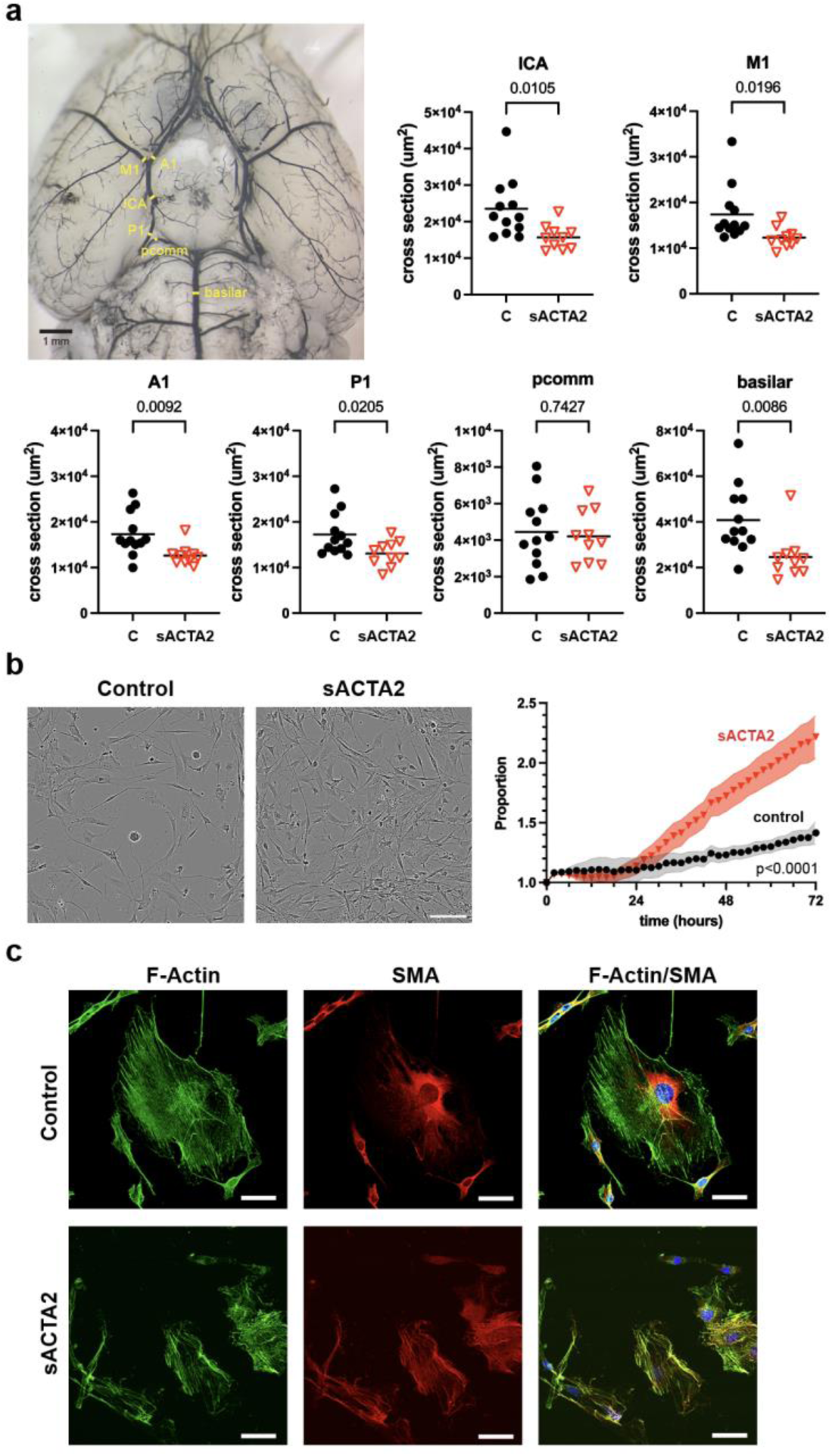
ACTA2 mice have vessel stenosis and increased proliferation of cells within the vessel wall. (a) Cross-sectional areas are smaller in many arteries within the circle of Willis in mice that express the cerebral and systemic ACTA2 R179H mutation (sACTA2) compared to control (C). Vessels are visualized after black ink injection and areas calculated from measured diameters at the internal carotid artery (ICA), first segment of the middle cerebral artery (M1), first segment of the anterior cerebral artery (A1), first segment of the posterior cerebral artery (P1), posterior communicating artery (pcomm), and basilar artery. (b) Cultured vascular smooth muscle cells from sACTA2 cerebral vessels proliferate at a greater rate than control (n = 9 per group). Scale bar 100 um. (c) F-actin (green) and smooth muscle actin (SMA, red) staining in control and sACTA2 cells. Control cells show organized stress fibers, whereas sACTA2 cells exhibit progressively distorted F-actin structure and abnormal SMA localization. Scale bar 50 um.

To assess whether these cellular abnormalities were associated with altered smooth muscle differentiation *in vivo*, we examined the contractile phenotype of cerebral vascular smooth muscle. Immunohistochemical analysis demonstrated reduced expression of SMA and smoothelin in cerebral arteries from ACTA2 mutant mice (**Figure 4a,b**). Transgelin and calponin showed similar downward trends that did not reach the threshold for statistical significance.

**Figure 4.**
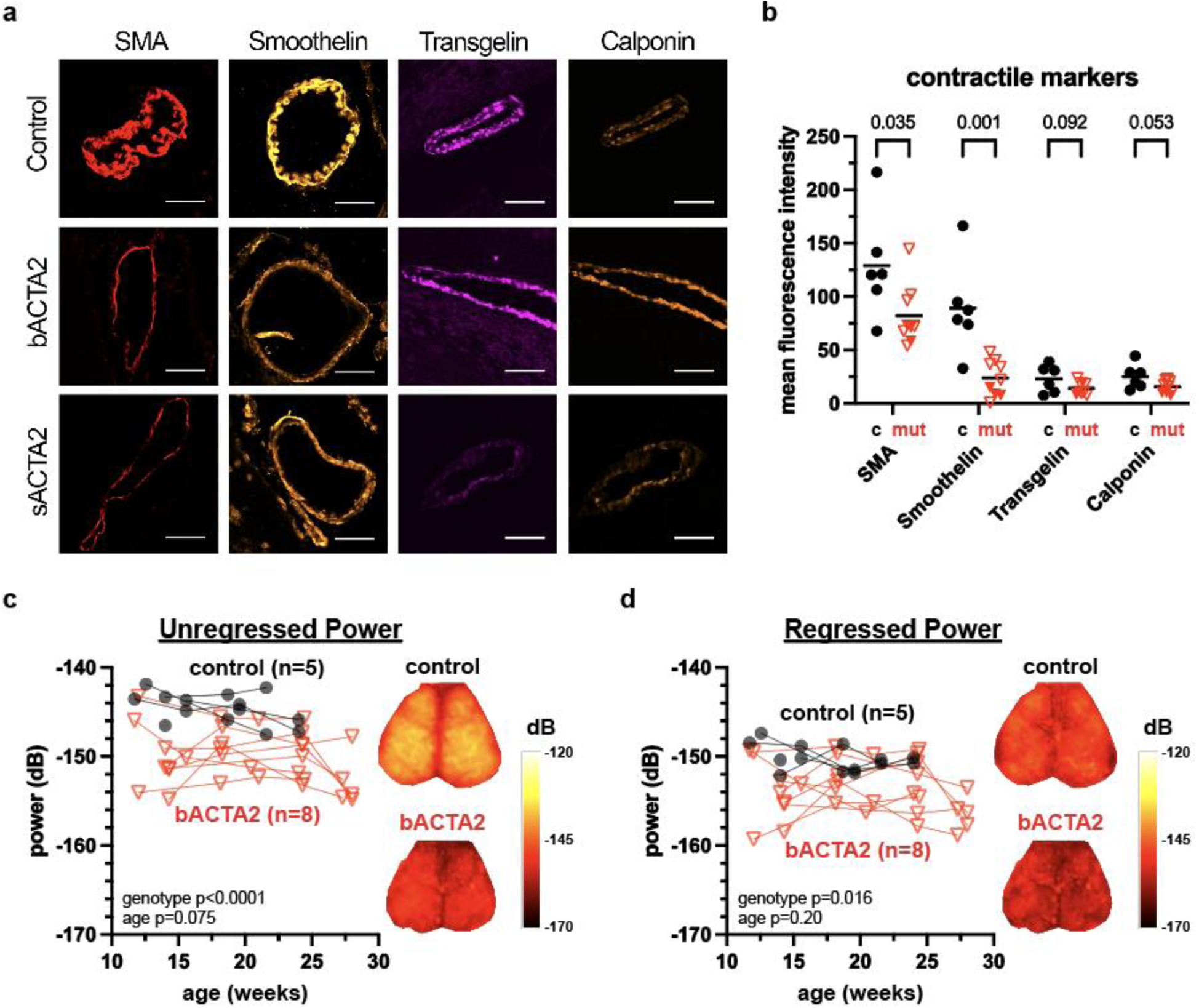
Contractile marker downregulation and attenuated spontaneous vascular fluctuations in ACTA2 mutant mice. (a) Representative cerebral artery cross sections processed with antibodies against contractile markers smooth muscle actin (SMA), smoothelin, transgelin, and calponin in control, brain and systemically expressed ACTA2 R179H (sACTA2), and brain specific expression of ACTA2 R179H (bACTA2). (b) Quantification of contractile marker mean fluorescence intensity in control (n = 6 circles), sACTA2 (n = 3 closed triangles), and bACTA2 (n = 6 open triangles) conditions. (c,d) Quantified total hemoglobin (HbT) fluctuations over the dorsal brain surface using longitudinal optical intrinsic signal imaging on a decibel power scale (dB) in control (n = 5) and bACTA2 (n = 8) mice. (c) Representative maps of unregressed HbT fluctuation power across the dorsal surface of the brain and a plot of the average power over time for control and bACTA2 mice, measured longitudinally as the mice age. (d) Representative power maps and plot of mean signal regressed HbT fluctuations.

Dynamic modulation of vascular tone is a key determinant of cerebrovascular reserve. Spontaneous cerebral blood-volume (CBV) fluctuations at rest were markedly reduced in mutant mice compared with controls (**Figure 4c,d**).^16^ This attenuation persisted whether all physiological fluctuations were included (**Figure 4c**) or physiological noise was regressed to enrich for neuronally driven hemodynamic oscillations analogous to the BOLD signal (**Figure 4d**).^9,17-19^

Systemic and cerebrovascular responses to vasoactive stimuli were markedly blunted in ACTA2 mutant mice (**Figure 5a–f**). Isoflurane anesthesia, which typically lowers blood pressure through vasodilation,^20^ was associated with higher resting arterial pressure in mutants compared with controls (**Figure 5b**), and this difference persisted under Avertin anesthesia (**Figure 5c**). Dose-response testing revealed attenuated hypotensive and hypertensive responses to sodium nitroprusside (SNP) and phenylephrine (PE), respectively (**Figure 5d,e**). Simultaneous laser speckle flowmetry demonstrated persistently lower cerebral blood flow (CBF) across systemic arterial pressures (**Figure 5f**).

**Figure 5.**
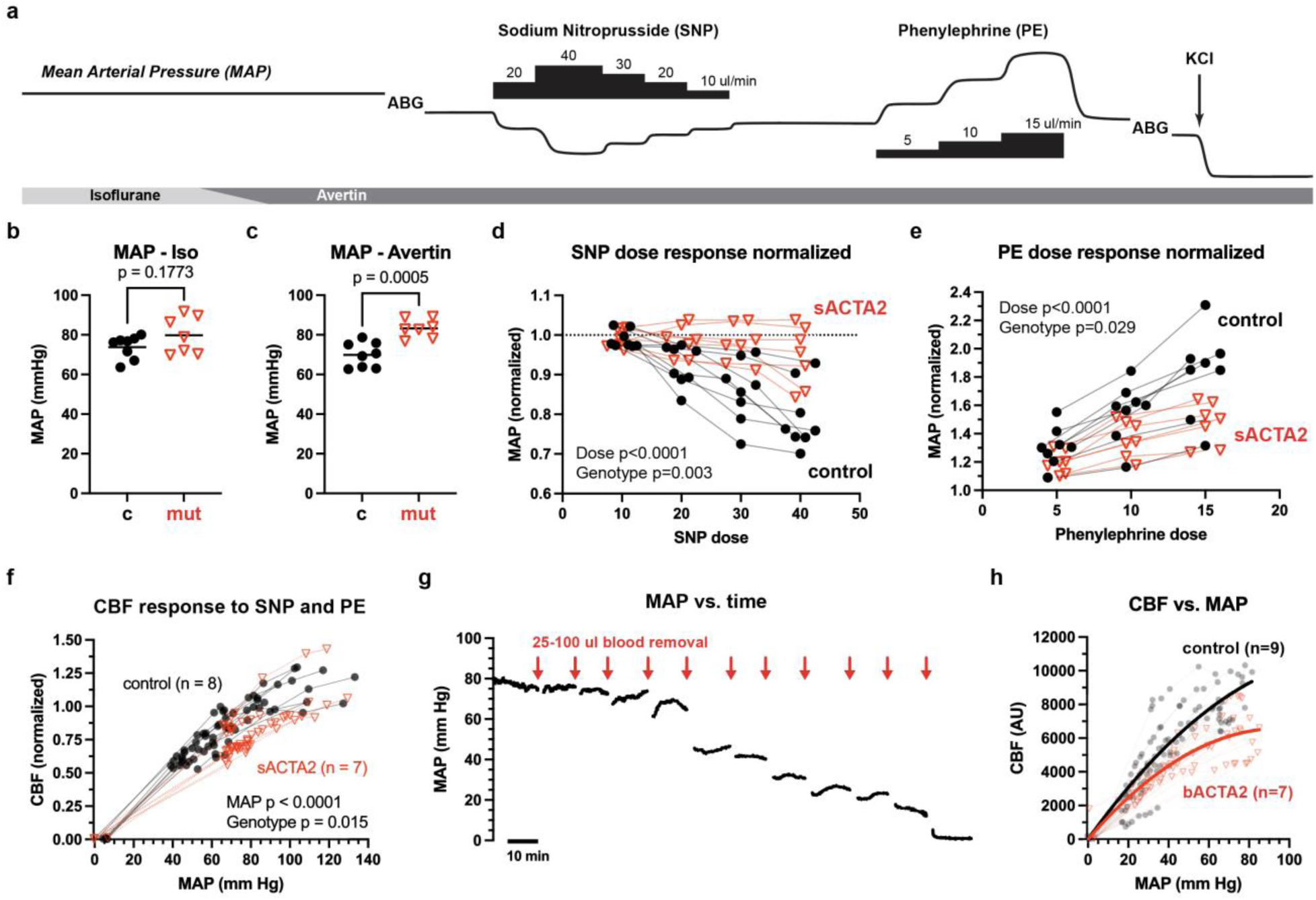
ACTA2 mice have impaired responsiveness to vasopressors and vasodilating agents and have lower cerebral blood flow (CBF). (a) Schematic of experimental timeline. (b-e) Systemic invasive mean arterial pressure (MAP) measurements under (b) isoflurane and (c) Avertin anesthetics, (d) the vasodilator sodium nitroprusside (SNP), and (e) the vasopressor phenylephrine (PE) in brain and systemically expressed ACTA2 R179H (sACTA2) mice. (f) The CBF vs. SNP- and PE-modulated MAP response in control vs. sACTA2 mice (n = 8 control, n = 7 sACTA2). (g) Blood pressure response to controlled exsanguination in a representative mouse expressing the ACTA2 R179H mutation specifically in the brain (bACTA). Mean arterial pressure (MAP) is determined with a catheter inserted into the femoral artery. Each red arrow represents removal of 25-100 ul blood through the same femoral arterial line. (h) The CBF vs. MAP response in control vs. bACTA2 mice during controlled exsanguination. P<0.0001 for condition and p<0.0001 for MAP when data are fit with a mixed effects model. Data from all mice and collected datapoints are shown (n = 9 control, n = 7 bACTA2). The best fit for each condition are shown as solid lines.

During controlled exsanguination, mean arterial pressure was reduced stepwise while CBF was measured over the dorsal cortex (**Figure 5g**). Mutant mice maintained lower CBF across the full pressure range (**Figure 5h**), revealing a downward-shifted pressure-flow relationship consistent with reduced cerebrovascular reserve.

Unilateral common carotid artery occlusion revealed impaired compensatory perfusion in ACTA2 mutant mice (**Figure 6**). Residual CBF following CCAO was lower in mutants compared with controls (**Figure 6c**). This physiological deficit was accompanied by increased mortality, greater weight loss, and worse behavioral outcomes (**Figure 6d-f**).

**Figure 6.**
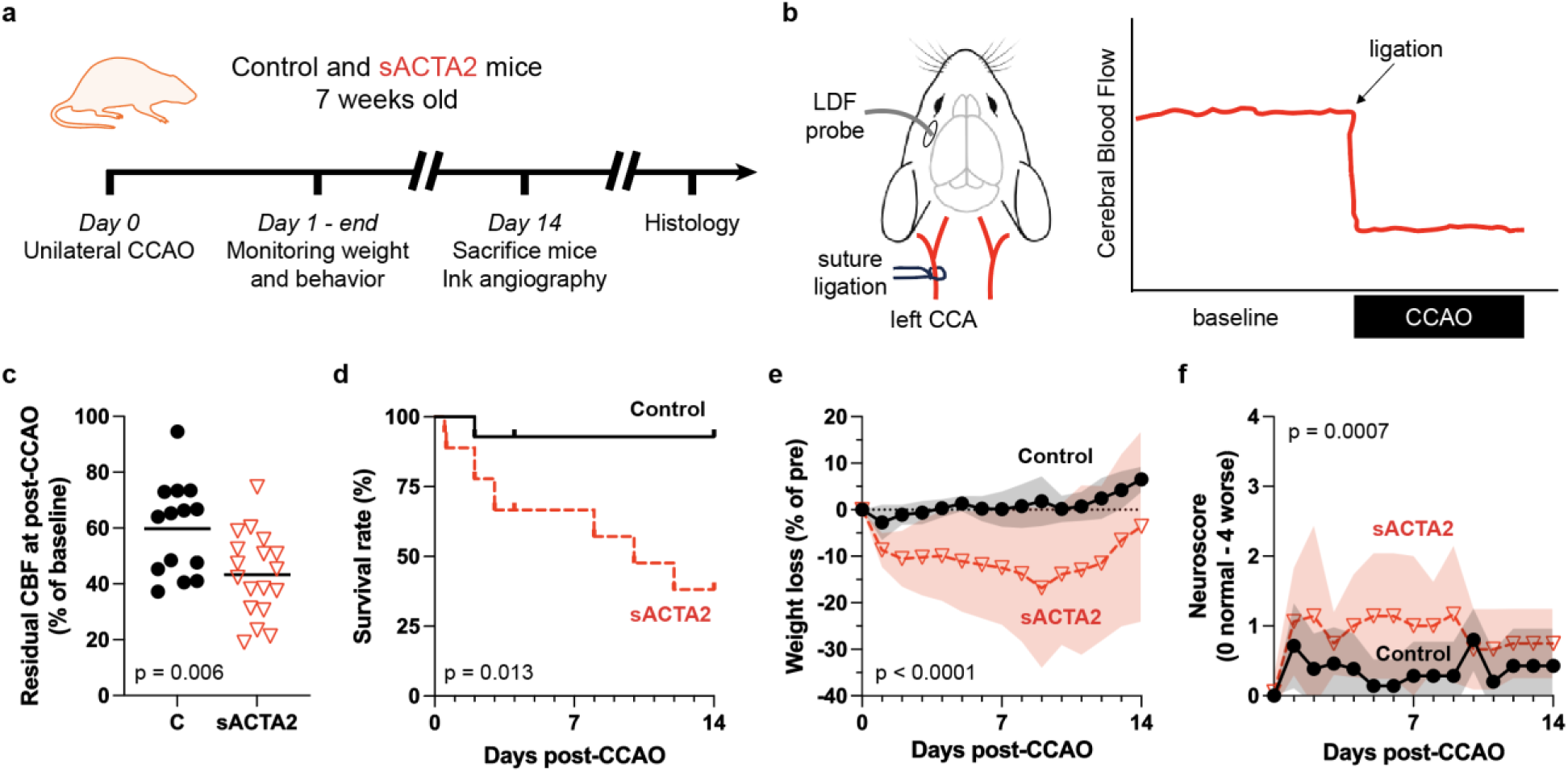
ACTA2 mutant mice exhibit an impaired ability to maintain CBF homeostasis under stress conditions. (a) Experimental timeline for controls and mice expressing the ACTA2 R179H mutation systemically and in the brain (sACTA2) that underwent permanent, unilateral common carotid artery occlusion (CCAO). (b) Illustration and schematic for the determination of relative CBF response to common carotid artery occlusion (CCAO) using laser Doppler flowmetry (LDF). (c) sACTA2 mice have lower residual CBF, (d) worse survival, (e) greater weight loss, and (f) worse neuroscores following ipsilateral CCAO (n = 14 control, n = 18 sACTA2).

Patients with ACTA2 R179H mutation exhibit early brain atrophy and white-matter injury.^13^ In the mouse model, white-matter staining intensity was reduced, with a qualitatively thinner corpus callosum and focal cavitations observed in a subset of ACTA2 mutants but not controls (**Figure 7a**). Luxol fast blue staining demonstrated reduced myelination in the corpus callosum, thalamus, and striatum (**Figure 7b**), with concordant reductions in myelin basic protein (MBP) immunostaining (**Figure 7c**). NeuN immunostaining revealed reduced cortical neuronal density in mutant mice (**Figure 7d**). To determine whether this structural brain injury was accompanied by functional consequences consistent with impaired perfusion control, we next assessed network-level brain function and behavior as downstream readouts of cerebrovascular reserve failure.

**Figure 7.**
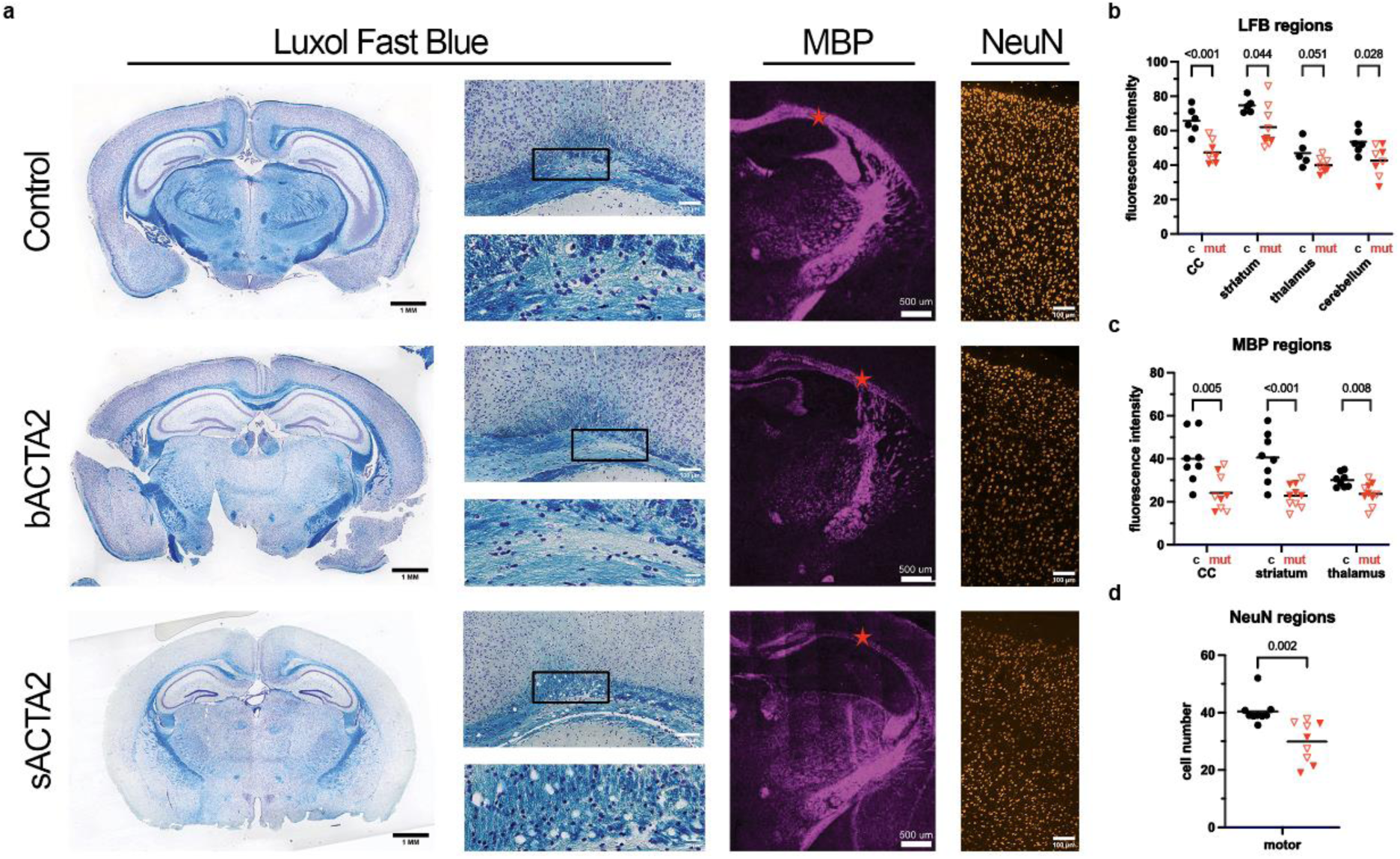
ACTA2 mice exhibit reduced myelination in white matter tracts and fewer cortical neurons. (a) Representative control, brain-specific ACTA2 R179H mice (bACTA2), and brain and systemically expressed ACTA2 R179H mice (sACTA2) coronal sections processed with luxol fast blue (LFB), anti-myelin basic protein (MBP) corpus coliseum marked with red star, and anti-NeuN (motor cortex). (b-d) Regional quantification of LFB, MBP, and NeuN in control and ACTA mutant (mut) mice. (b) LFB fluorescence intensity in corpus callosum (CC), striatum, thalamus, and cerebellum in ACTA2 mutant (bACTA2 n = 6, sACTA2 n = 4) and control mice (n = 6). (c) MBP fluorescence intensity in CC, striatum, and thalamus in ACTA2 mutant (bACTA2 n = 5, sACTA2 n = 5) and control mice (n = 8). (d) Number of cells expressing NeuN demonstrates lower density of neurons in ACTA2 mutant (bACTA2 n = 5, sACTA2 n = 4) compared to control mice (n = 9). bACTA2 (open triangles) and sACTA2 (closed triangles).

Resting state functional connectivity was impaired in mice expressing the brain-specific ACTA2 R179H mutation (**Figure 8a**). Optical intrinsic signal imaging^21^ revealed reduced interhemispheric and global connectivity, most prominently in total hemoglobin (HbT) and oxyhemoglobin (HbO) signals. Connectivity declined further with age when assessed using HbT-based metrics, whereas age-related effects were not observed in HbO- or HbR-based measures (**Figure 8a**). Behavioral testing in bACTA2 mice revealed functional consequences of cerebrovascular reserve failure. Despite preserved gross neurological function, mutant mice exhibited reduced ambulation, increased anxiety-like behavior, and impaired exploratory and working memory performance (**Figure 8b-g**).

**Figure 8.**
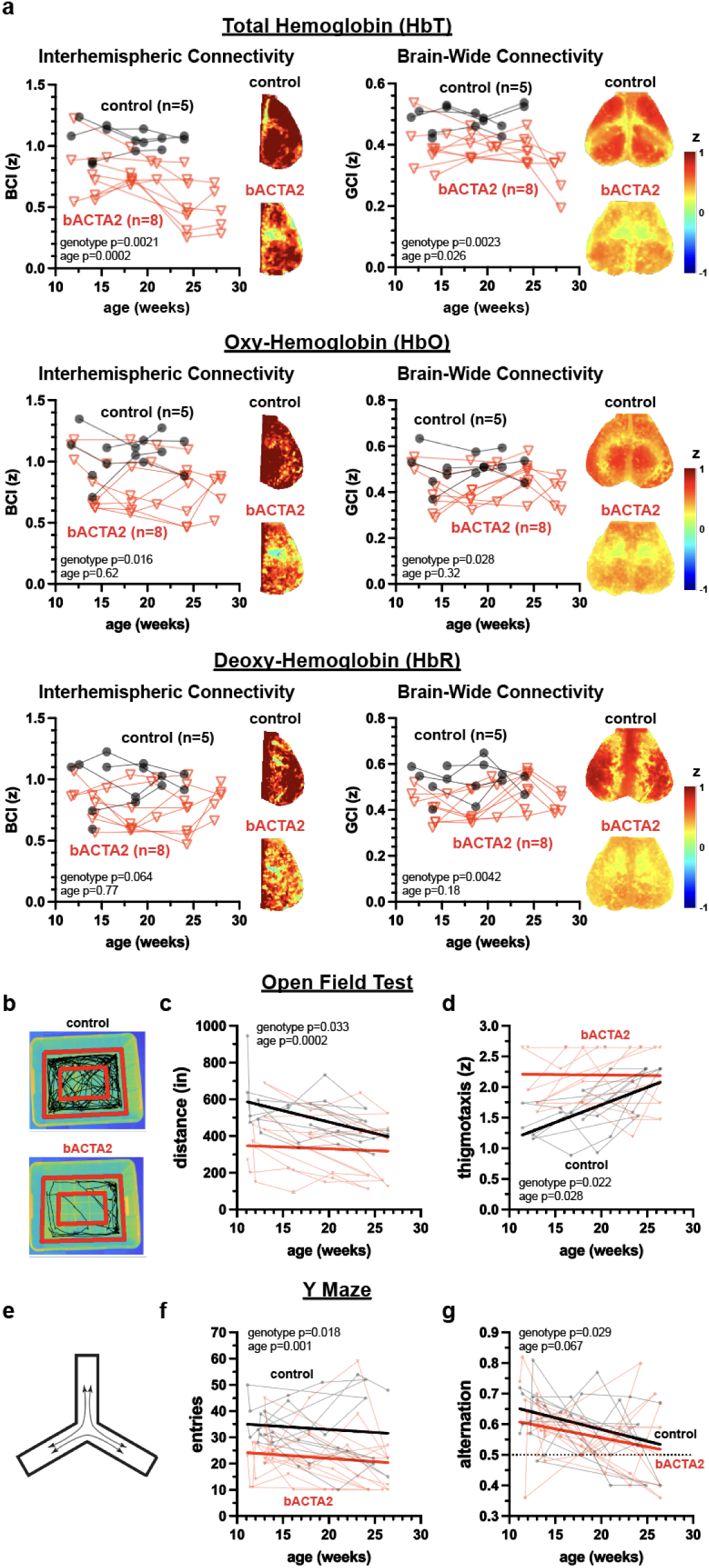
Cerebrovascular reserve failure is associated with network-level dysfunction and behavioral impairment. (a) ACTA2 mutant mice exhibit reduced resting state functional connectivity, consistent with impaired neurovascular coupling downstream of cerebrovascular reserve failure. Fisher z-transformed bihemispheric connectivity indices (BCI) quantify interhemispheric functional connectivity between homotopic cortical regions in control and brain-specific ACTA2 R179H (bACTA2) mice. Representative maps depict reduced interhemispheric connectivity in bACTA2 mice. Global connectivity indices (GCI) quantify brain-wide network integration. Connected lines denote longitudinal measurements within individual animals. Connectivity analyses based on total hemoglobin (HbT), oxyhemoglobin (HbO), and deoxyhemoglobin (HbR) reveal signal-dependent effects reflecting their differential sensitivity to vascular and neuronal contributions. (b-g) Behavioral consequences of impaired cerebrovascular reserve and network-level dysfunction. (b) Representative open-field test (OFT) movement trajectories in control and bACTA2 mice. (c) Longitudinal ambulation distance over 5 minutes. (d) Fisher z-transformed thigmotaxis scores reflecting anxiety-like behavior. (e) Schematic of the Y-maze spontaneous alternation task. (f) Total number of arm entries over 8 minutes. (g) Fraction of spontaneous alternations, indicating impaired spatial working memory in bACTA2 mice.

## Discussion

In this study, we identify baseline impairment of cerebrovascular reserve and autoregulation as a central physiological consequence of pathogenic smooth muscle dysfunction in ACTA2 R179H-associated multisystemic smooth muscle dysfunction syndrome (MSMDS). Using a genetically defined mouse model, we demonstrate a convergent set of vascular abnormalities, including arterial rectification and narrowing, impaired vasodynamic activity, blunted vasoreactivity, and a downward shift in the blood pressure-cerebral blood flow relationship. Together, these abnormalities limit the ability of the cerebral circulation to maintain perfusion across physiologic stressors. As a result, ACTA2 mutant mice exhibit reduced tolerance to hypotension and arterial obstruction, mirroring the clinical vulnerability to watershed ischemia observed in MSMDS patients during anesthesia, systemic illness, or dehydration.^13^

The downstream brain phenotype observed in ACTA2 mutant mice supports a vascular-first mechanism in which impaired vascular adaptability and perfusion control drive structural and functional decline. Mutant mice exhibited reduced white matter integrity, neuronal loss, impaired functional connectivity, and behavioral deficits that emerged in parallel with cerebrovascular reserve failure. These findings point to impaired cerebrovascular reserve and neurovascular coupling as the primary drivers of network and behavioral dysfunction. This pattern is consistent with a model in which chronic and stress-induced hypoperfusion promotes end-organ injury, particularly within white matter networks that are highly sensitive to modest or intermittent reductions in blood flow. Establishing causality and reversibility will require targeted rescue experiments.

These findings place ACTA2 vasculopathy within a broader framework implicating vascular dysfunction as a primary contributor to cognitive impairment and neurodegeneration.^1-4^ While vascular abnormalities are often considered secondary modifiers in disorders such as Alzheimer’s disease, ACTA2 R179H represents a human experiment of nature in which the initiating insult is purely vascular. The reduced spontaneous hemodynamic fluctuations observed in ACTA2 mutant mice parallel abnormalities reported in cerebral small vessel disease, aging, and dementia,^22,23^ reinforcing the importance of vascular tone dynamics and reserve capacity as unifying mechanisms across diverse forms of vascular cognitive impairment.

Our findings complement and extend a recent report describing rescue of systemic features of ACTA2 R179H disease using genomic editing in a constitutive knock-in mouse model.^24^ That study reported lower systemic blood pressure in mutant animals, whereas we observed higher arterial pressures under anesthesia. These differences likely reflect methodological context: blood pressure in the prior study was measured non-invasively in awake animals using tail-cuff plethysmography, whereas our measurements were obtained invasively under isoflurane anesthesia, a condition that normally induces vasodilation. Under these conditions, higher pressure in ACTA2 mutant mice is most parsimoniously interpreted as impaired vasodilatory capacity, consistent with our demonstration of blunted responses to vasoactive agents and reduced cerebrovascular reserve.

A complementary recent study by Kaw et al. used an SMC-targeted Acta2^R179C/+^ mouse model to implicate incomplete smooth muscle differentiation in the development of moyamoya-like occlusive lesions.^25^ In that work, smooth muscle cells exhibited metabolic reprogramming characterized by reduced oxidative phosphorylation and increased glycolytic traits. Notably, because their mice lacked overt distal internal carotid occlusions or brain ischemia at baseline, carotid ligation was required to provoke occlusive lesions and neurologic injury, which were prevented by nicotinamide riboside treatment. This injury-provoked model complements our findings by highlighting an occlusive remodeling pathway. In parallel, our study defines a baseline physiological phenotype (i.e., impaired vasodynamics and reduced cerebrovascular reserve) that is detectable without vascular injury and manifests clinically under stress.

Together, these studies support a dual-mechanism framework for ACTA2-associated cerebrovascular disease. Impaired smooth muscle differentiation and maladaptive phenotypic switching can promote occlusive remodeling following vascular injury, while baseline impairment of vascular tone regulation and reserve capacity creates vulnerability to hypoperfusion and watershed ischemia even in the absence of fixed occlusion. Both mechanisms are likely to contribute to disease expression in patients, with their relative importance shaped by developmental context, systemic stressors, and vascular injury.

Our functional connectivity analyses further support a vascular origin of network dysfunction. Connectivity reductions were most robust in hemoglobin signals closely linked to neuronal activity and neurovascular coupling, particularly HbO and HbT.^26-28^ In contrast, HbR-based measures—analogous to the BOLD signal used in human fMRI—were less sensitive, underscoring inherent limitations of clinical imaging in resolving subtle neurovascular deficits.^29^ The ability to interrogate these signals in animal models provides mechanistic insight into how impaired vascular reserve and tone regulation disrupt large-scale brain networks.

Several limitations warrant consideration. Rescue experiments using vascular gene therapy or pharmacologic strategies aimed at restoring vascular reserve will be required to further establish causality and reversibility of end-organ brain injury. Species-specific differences in cerebrovascular architecture may influence disease expression, and early mortality in sACTA2 mice may introduce survival bias, potentially underestimating the full impact of smooth muscle dysfunction on cerebrovascular physiology.

In summary, we establish a robust model in which genetically defined smooth muscle dysfunction is associated with baseline impairment of cerebrovascular reserve, rendering the brain vulnerable to hypoperfusion and stress-induced ischemic injury. By linking impaired vascular tone regulation and autoregulation to network disruption and behavioral dysfunction, this work defines a physiological mechanism that operates independently of fixed arterial occlusion. More broadly, these findings highlight cerebrovascular reserve as a central physiological axis linking systemic vascular dysfunction to long-term brain health. Together with recent studies demonstrating injury-provoked occlusive remodeling in ACTA2 vasculopathy, our findings highlight cerebrovascular reserve as a critical therapeutic target in MSMDS and more broadly in hypotension-prone vasculopathies and conditions characterized by impaired vascular reactivity.

## Author Contributions

Takahiko Imai – Conceptualization; Methodology; Investigation (laser speckle CBF, blood pressure/vasoreactivity, black ink angiography, CCAO); Formal Analysis; Visualization; Writing—Original Draft; Writing—Review & Editing.

Vijai Krishnan – Conceptualization; Methodology; Investigation (MRI/MRA); Formal Analysis; Visualization; Project Administration; Writing—Review & Editing.

James H. Lai – Methodology; Investigation (optical intrinsic signal imaging / resting state functional connectivity and behavioral assays); Software; Formal Analysis; Visualization; Writing—Review & Editing.

Elyssa Alber – Investigation (laser speckle CBF support, behavioral assays, genotyping); Data Curation; Writing—Review & Editing.

Lydia Hawley – Investigation (MRI support, behavioral assays, genotyping); Data Curation; Writing—Review & Editing.

Pazhanichamy Kalailingam – Methodology; Investigation (histology and immunostaining, including CCAO-related histology); Formal Analysis; Writing—Review & Editing.

Aarushi Gandhi – Investigation (histology imaging); Formal Analysis (histology quantification support); Visualization support; Writing—Review & Editing.

Joanna Yang – Methodology; Investigation (laser speckle CBF, blood pressure/vasoreactivity, MRI, behavioral assays); Software; Formal Analysis; Visualization; Writing—Review & Editing.

Diana Tambala – Investigation (human neuroimaging curation and clinical coordination); Visualization (clinical figure preparation); Writing—Review & Editing.

David C. Hike – Investigation (MRI acquisition); Data Curation; Writing—Review & Editing.

Xiaoqing Alice Zhou – Software; Formal Analysis (MRI/MRA processing and analysis); Visualization support; Writing—Review & Editing.

Claire Fong – Investigation (genotyping and colony coordination; histology imaging support); Data Curation; Writing—Review & Editing.

Benjamin Ondeck – Investigation (patient recruitment and data collection); Data Curation; Writing—Review & Editing.

Emily T Da Cruz – Investigation (patient recruitment and data collection); Data Curation; Writing—Review & Editing.

Xiaochen Liu – Software; Formal Analysis (MRI/MRA processing and analysis); Writing—Review & Editing.

Angelyna K. Siv – Investigation (cranial window preparation and behavioral assays); Writing—Review & Editing.

Miran Öncel – Investigation (resting state imaging); Formal Analysis support; Writing—Review & Editing.

Sabyasachi Das – Resources; Writing—Review & Editing.

Sava Sakadžić – Conceptualization; Methodology (optical intrinsic signal imaging and CBV fluctuation analysis); Supervision; Writing—Review & Editing.

Cenk Ayata – Conceptualization; Methodology (cerebrovascular physiology); Supervision; Writing—Review & Editing.

Xin Yu – Conceptualization; Methodology (MRI/MRA); Supervision; Writing—Review & Editing.

Mark E. Lindsay – Conceptualization; Supervision; Resources; Funding Acquisition; Writing—Review & Editing.

Patricia L. Musolino – Conceptualization; Supervision; Resources; Funding Acquisition; Writing—Review & Editing.

David Y. Chung – Conceptualization; Methodology; Investigation; Formal Analysis; Visualization; Writing—Original Draft; Writing—Review & Editing; Supervision; Project Administration; Resources; Funding Acquisition.

## Acknowledgements

This work was supported by the National Institutes of Health (R01NS125353 to M.E.L. and P.M., K08NS112601 to D.Y.C., and R01NS136224 to D.Y.C.). Portions of this manuscript were edited and refined with the assistance of an artificial intelligence-based large language model to improve clarity, organization, and readability of the text. The AI tool was not used to generate original scientific content, analyze data, interpret results, or create figures. All authors reviewed, edited, and approved the final manuscript and assume full responsibility for the accuracy and integrity of the content.

